# Early hippocampal hyperexcitability followed by disinhibition in a mouse model of Dravet syndrome

**DOI:** 10.1101/790170

**Authors:** Yael Almog, Marina Brusel, Karen Anderson, Moran Rubinstein

## Abstract

Dravet syndrome (Dravet) epilepsy begins with febrile seizures followed by worsening to refractory seizures, with some improvement and stabilization toward adolescence. The neuronal basis of Dravet is debatable, with evidence favoring reduced inhibition or enhanced excitation. Focusing on the firing properties of hippocampal CA1 pyramidal neurons and oriens-lacunosum moleculare (O-LM) interneurons, we provide a comprehensive analysis of the activity of both cell types through the febrile, worsening and stabilization stages. Our data indicate a temporary increase in the excitability of CA1 pyramidal neurons during the febrile stage, which is fully reversed by the onset of spontaneous seizures. In contrast, reduced function of O-LM interneurons persisted from the febrile through the stabilization stages, with the greatest impairment of excitability occurring during the worsening stage. Thus, both excitatory and inhibitory neurons contribute to Dravet, indicating complex and reciprocal pathophysiological neuronal changes during the different stages of the disease.

**Graphical abstract:** 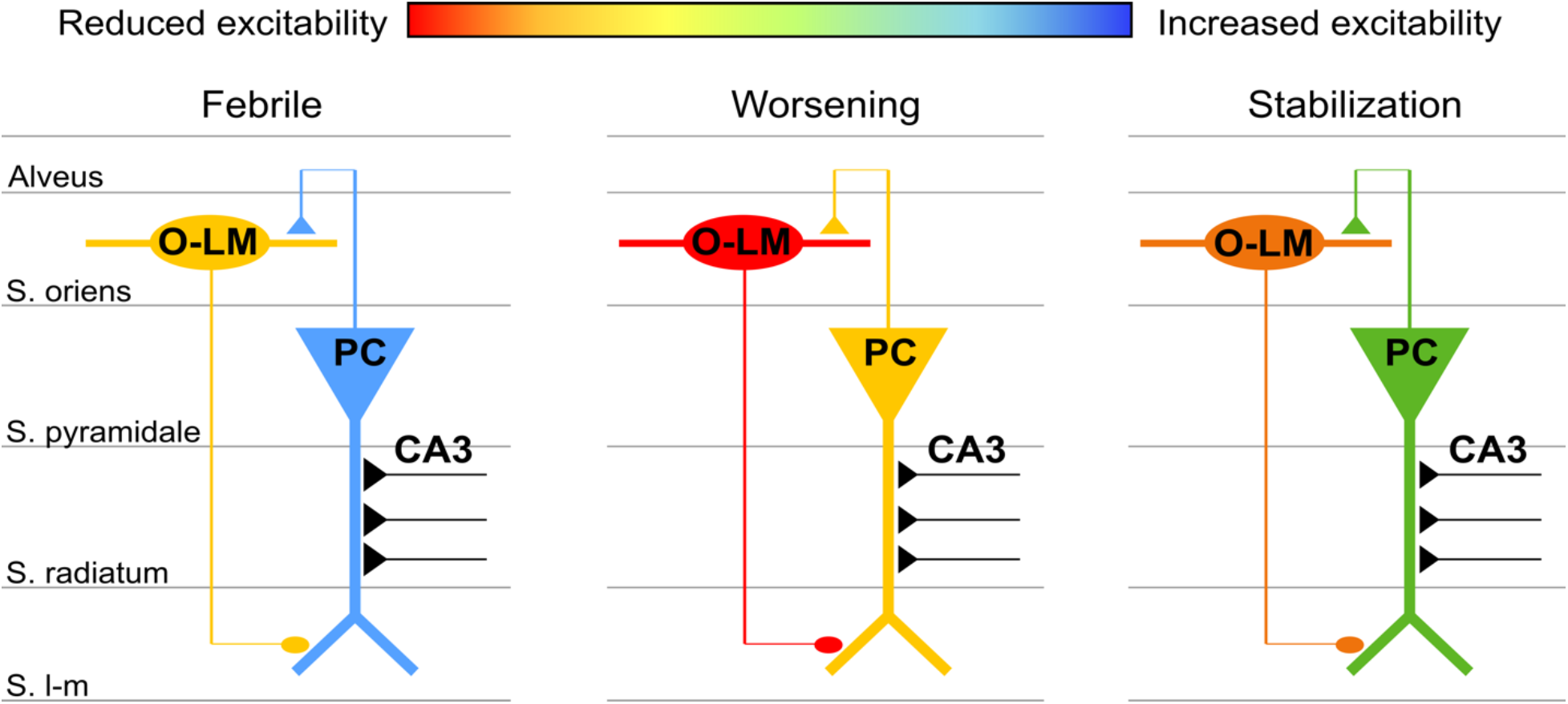

## Introduction

Dravet syndrome (Dravet) is a rare, severe childhood onset epilepsy. Most Dravet cases arise from heterozygous *de novo* mutations in the *SCN1A* gene (Claes et al., 2009), which encodes for the alpha subunit of voltage gated sodium channel type I (Na_v_1.1). The disease first manifests with febrile seizures occurring around six months of age. This “febrile stage” progresses between the first and second years of life to the “worsening stage,” which is characterized by spontaneous refractory seizures and frequent episodes of status epilepticus, as well as global developmental delay. Around age six, the epileptic phenotypes improve and seizure burden is reduced. However, the global developmental delay persists in this “stabilization stage” (Dravet, 2011; Dravet and Oguni, 2013; Gataullina and Dulac, 2017).

Murine models of Dravet (DS mice), presenting with severe epilepsy, cognitive and social deficits, recapitulate the human disease (Han et al., 2012; Ito et al., 2013; Kalume et al., 2013; Ogiwara et al., 2007; Yu et al., 2006). Despite extensive studies of these models, the neuronal basis of Dravet and the mechanisms underlying the different stages are still debated, with evidence in favor of reduced function of inhibitory neurons, or enhanced function of excitatory neurons. In support of the “inhibitory neuron” hypothesis, electrophysiological studies demonstrated impaired firing of multiple types of inhibitory neurons during the worsening stage, with no apparent change in the excitability of excitatory neurons (De Stasi et al., 2016; Favero et al., 2018; Goff and Goldberg, 2019; Han et al., 2012; Ogiwara et al., 2007; Rubinstein et al., 2015b, 2015a; Tai et al., 2014; Tsai et al., 2015). Moreover, selective deletion of *Scn1a* in inhibitory neurons was sufficient to cause seizures and premature mortality (Cheah et al., 2012; Kalume et al., 2013; Kuo et al., 2019; Ogiwara et al., 2013; Rubinstein et al., 2015a). Conversely, inhibitory neurons of the reticular thalamic nucleus were hyperexcitable (Ritter-Makinson et al., 2019), and reduced function of cortical parvalbumin (PV)-positive interneurons was shown to be limited only to the worsening stage (Favero et al., 2018). Moreover, other studies implicated the involvement of excitatory neurons, demonstrating increased sodium conductance in hippocampal dissociated glutamatergic neurons of DS mice and patient-derived neurons (Liu et al., 2013; Mistry et al., 2014).

Here, using the commercially available DS mice that harbor a missense *Scn1a^A1783V^* mutation, we examined the firing properties of hippocampal CA1 pyramidal neurons and stratum oriens-lacunosum moleculare (O-LM) inhibitory interneurons at the three different stages of Dravet. By focusing on responses to synaptic stimulation, we show a temporary increase in the excitability of CA1 pyramidal neurons that was limited to the febrile stage. This transient hyperactivity was fully reversed by the worsening stage, with complete convalesce at the stabilization stage. In contrast, impairment in the function of O-LM interneurons was evident from the febrile through the stabilization stages, with the greatest dysfunction occurring at the worsening stage. These data show a biphasic pathophysiological mechanism, beginning with the contribution of excitatory neurons to the initial stage of Dravet and followed by disinhibition. This unexpected time course indicates that complex and reciprocal neuronal changes govern the developmental trajectory of the disease.

## Results

### Susceptibility to thermally induced seizures correlates with the severity of epilepsy

DS mice harboring the *Scn1a^A1783V^* missense mutation, maintained on the pure C57BL/6J genetic background, develop normally during their first weeks of life. Spontaneous seizures very rarely occurred before postnatal day (P) 16 but were frequently observed during routine handling in mice during their fourth week of life (P20-P28). The overall mortality of this cohort was ~50% (Fig. 1A). Death was observed after P18; with most of the deaths (86%) occurring between P20 and P28. Mice that survived beyond this fourth week still experienced spontaneous seizures, but the rate of death steeply declined, with 7.4% of the deaths observed between P29 and P33. This survival curve indicates that the febrile stage in DS mice ends around P18, the worsening stage corresponds to the fourth week of life (P20-P28) and the stabilization stage starts after P30.

**Figure 1.**
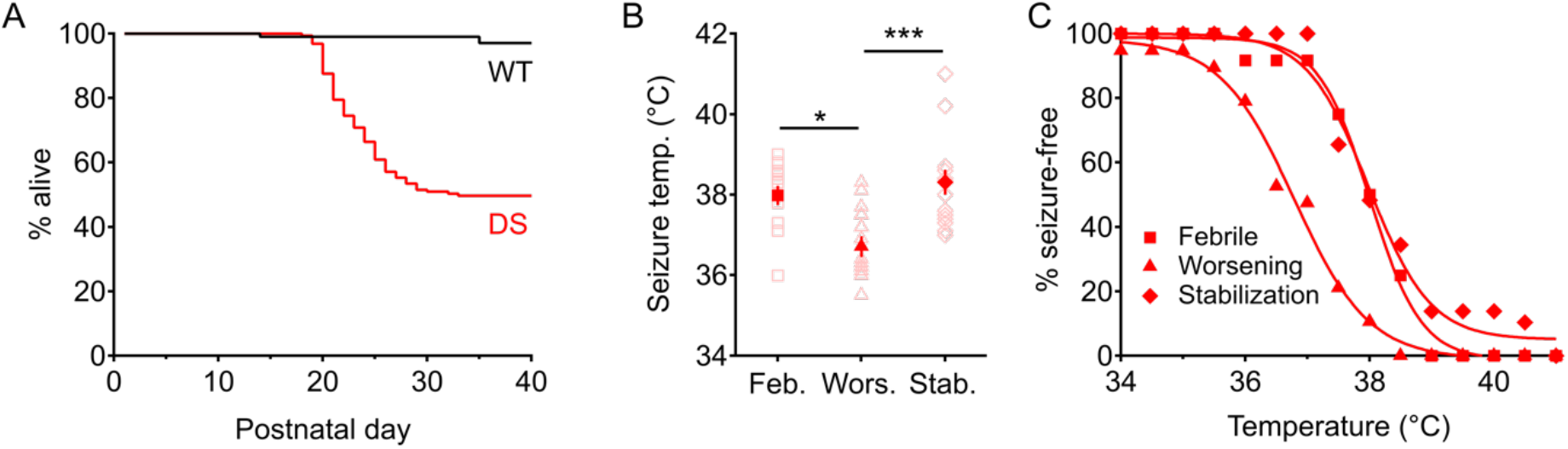
Mortality and susceptibility to thermally induced seizures in DS mice. (A) Survival curves of DS (n=161) and WT (n=102) mice. (B) Open symbols depict the distribution of seizure temperatures during the febrile stage (n=12), the worsening stage (n=19) and the stabilization stage (n=16). Filled red symbols indicate mean values ± SEM. WT mice had no seizures within these temperature range (P14: n=11; P21: n=11; P35: n=6). Statistical analysis utilized One-Way ANOVA, followed by Tukey *post-hoc* analysis. \**P* < 0.05; \*\*\**P* < 0.001. (C) Percentage of mice (the same data as in B) remaining free of thermally induced seizures at the indicated temperatures.

We next examined the sensitivity to thermal induction of seizures in mice of different ages. All of the tested DS mice had generalized tonic-clonic seizures at temperatures that are well within the range of childhood fever (Fig. 1B, C). At P14-P16, during the mostly quiescent febrile stage, DS mice had thermally induced seizures at- or below 39°C. A week later, during the worsening stage, seizures occurred at lower, near-normal body core temperature (36.72 ± 0.26 °C). By the stabilization stage, seizure temperature was again higher, and similar to that of mice during the febrile stage (Fig. 1B, C).

Together, these data confirm that DS mice carrying the *Scn1a^A1783V^* missense mutation recapitulate Dravet epilepsy similarly to earlier DS models which are based on *Scn1a* truncation mutations (Ogiwara et al., 2007; Yu et al., 2006). Moreover, these mice present the three stages of Dravet, with (*i*) the febrile stage preceding the onset of spontaneous seizures, (*ii*) the worsening stage characterized by increased mortality and high thermal sensitivity, and (*iii*) the stabilization stage exhibiting reduced mortality rate and lower susceptibility to thermal induction of seizures.

### Reduced intrinsic excitability of O-LM cells at the onset of epilepsy

There are conflicting results regarding the neuronal basis of Dravet, with evidence for reduced inhibition (Cheah et al., 2012; De Stasi et al., 2016; Ogiwara et al., 2007; Rubinstein et al., 2015b; Tai et al., 2014; Yu et al., 2006) that might be transient (Favero et al., 2018), as well as indications of increased activity of excitatory neurons (Liu et al., 2013; Mistry et al., 2014). Here, we focused on the hippocampal CA1 region, which has been shown to be important for Dravet epilepsy (Stein et al., 2019), examining the function of CA1 pyramidal neurons and stratum oriens interneurons (O-LM) at the three stages of Dravet. Within the CA1 microcircuit, recurrent collaterals of CA1 pyramidal cells activate O-LM interneurons, which in turn mediate feedback inhibition of these same excitatroy neurons (Klausberger and Somogyi, 2008; Müller and Remy, 2014; Scanziani and Pouille, 2004) (Fig. S1), controlling the final output of the hippocampus to extra-hippocampal regions.

Analysis of the firing properties of O-LM cells in response to injection of 1 s long depolarizing current through the patch pipette showed unaltered firing during the febrile stage, and reduction of the number of action potentials (APs) during the worsening and the stabilization stages (Fig. 2A-D). Examination of the firing close to the rheobase, which better resembles firing under physiological conditions, showed that while the firing impairment persists form the worsening to the stabilization stage, the severity of this impairment tends to decline (Fig. 2E-G).

**Figure 2.**
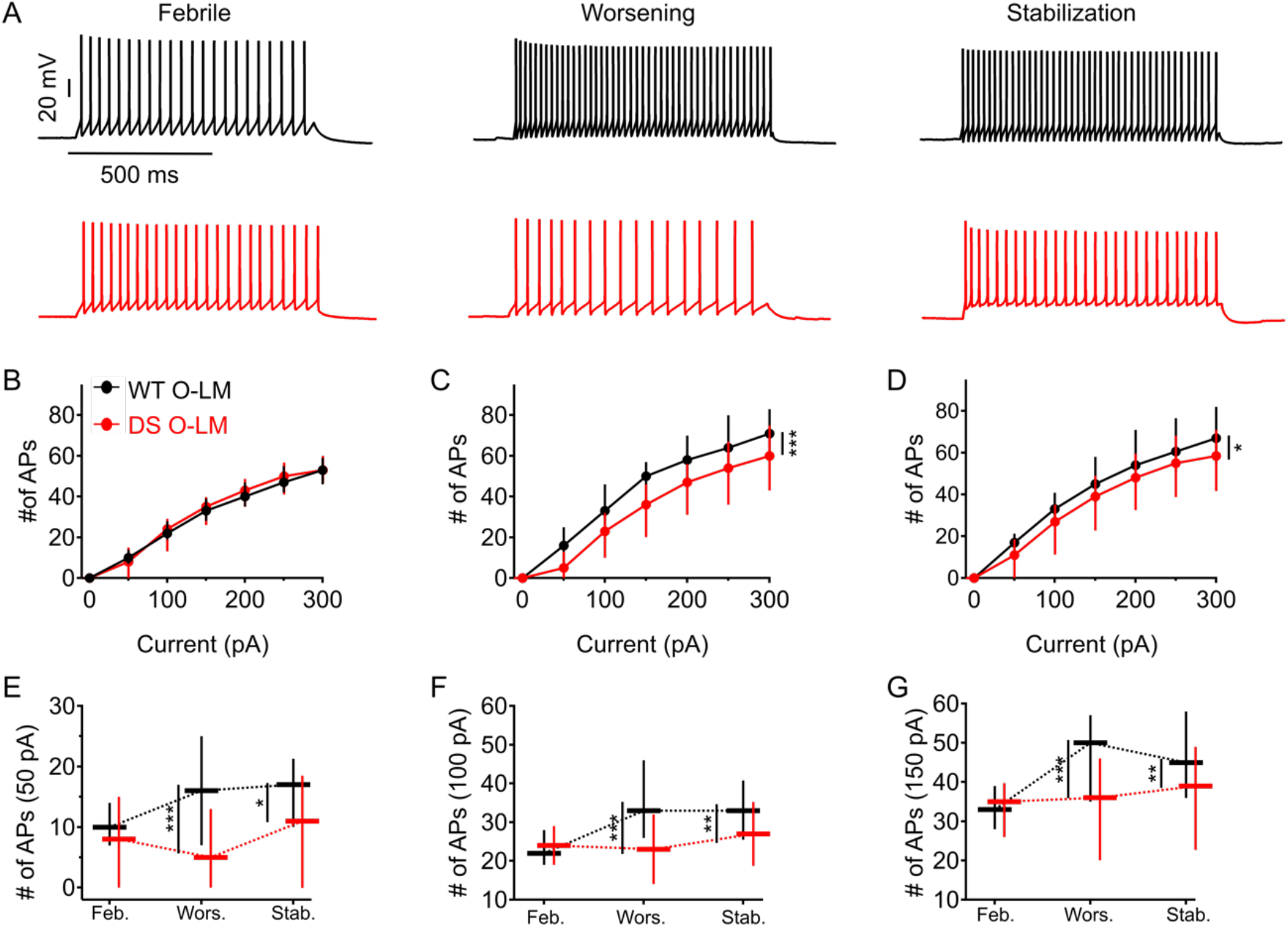
Reduced firing of O-LM interneurons during the worsening and stabilization stages. (A) Sample traces of whole-cell current clamp recordings from WT (black) and DS (red) O-LM interneurons in response to current injection of 100 pA through the patch pipette. (B-D) AP counts in response to 1 s depolarizing current injection (also corresponding to firing frequency), at the indicated intensities, during the febrile (B), worsening (C) and stabilization (D) stages. Symbols represent medians while vertical line segments under and above the medians represent the 25 and 75 percentile ranges, respectively. Statistical analysis utilized Two-Way Repeated Measures ANOVA, followed by Holm-Sidak *post-hoc* analysis. (E-G) AP counts generated in response to 50 pA (E), 100 pA (F), or 150 pA (G) in WT and DS O-LM neurons during the different stages of Dravet. Data are presented as median (horizontal midlines) and range (vertical line segments under or over the medians represent the 25 and 75 percentile ranges, respectively). Statistical analysis was done using Two-Way ANOVA, followed by Holm-Sidak *post-hoc* analysis. WT: n=39-41; DS: n=30-44 (see Table 1A for more details). \**P* < 0.05, \*\*\**P* < 0.001.

Examination of the properties of single APs showed no changes during the febrile stage, and increased rheobase and threshold with reduced input resistance at the worsening stage. At the stabilization stage, the rheobase and input resistance were corrected, but the threshold for AP remained increased (Fig. 3 and Table 1A). Thus, the properties of APs also indicate reduced function at the worsening stage that is partially corrected at the stabilization stage.

**Figure 3.**
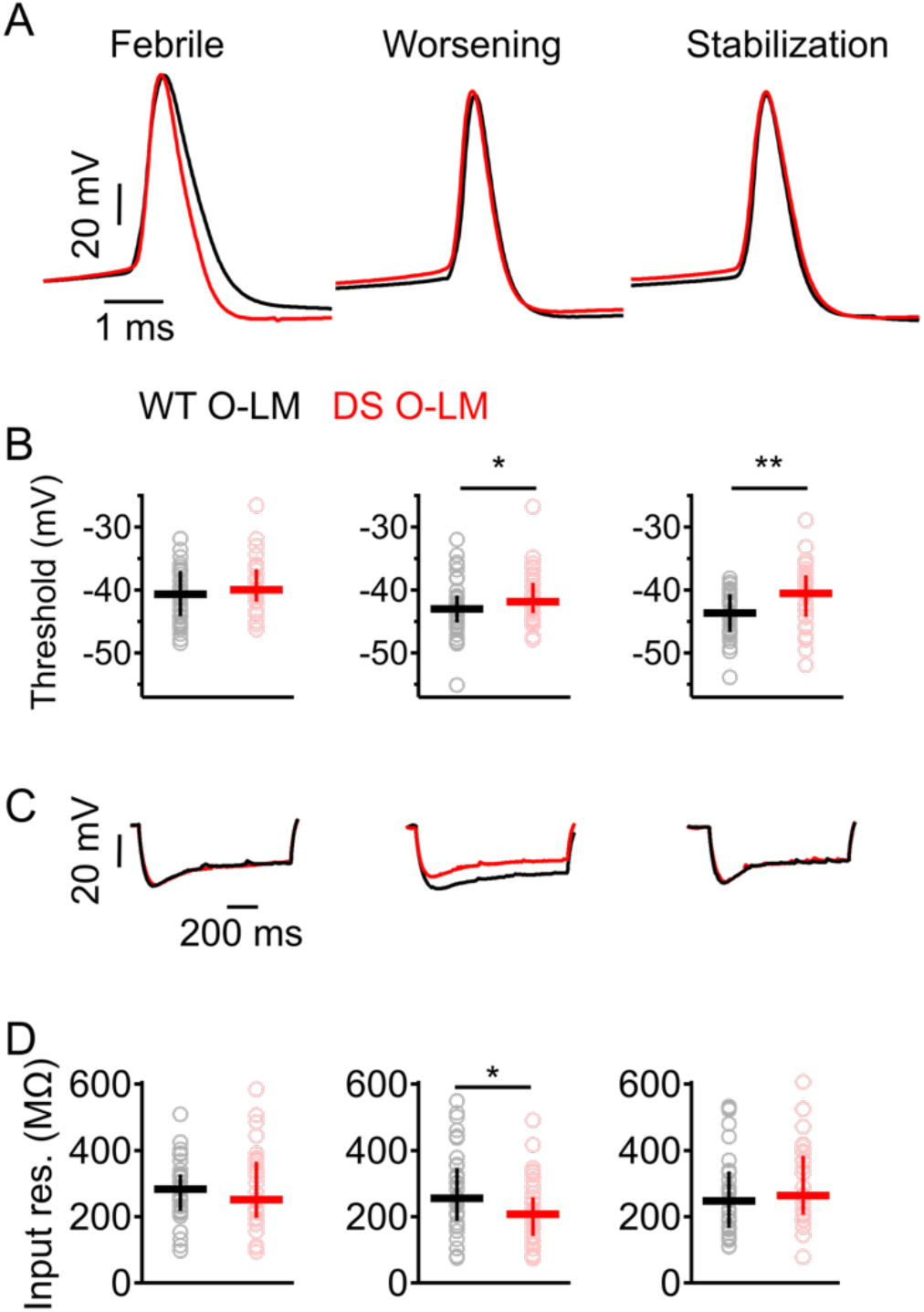
Increased threshold in DS O-LM interneurons during the worsening and stabilization stages. (A) Representative APs from WT (black) and DS (red) O-LM interneurons at different ages. Firing was induced by 10 ms depolarizing current injected through the patch pipette at rheobase. (B) AP threshold at the indicated Dravet stages. Statistical analysis was performed using Mann-Whitney Rank Sum Test. (C) Representative traces of voltage changes induced by injection of −100 pA. (D) Input resistance at the indicated ages. Data are presented as median (horizontal midlines) and range (vertical line segments under and over medians represent the 25 and 75 percentiles, respectively). Statistical analysis was performed using Mann-Whitney Rank Sum Test. WT: n=39-41; DS: n=30-44 (see Table 1A for more details). \**P* < 0.05, \*\**P* < 0.01.

**Table 1:**
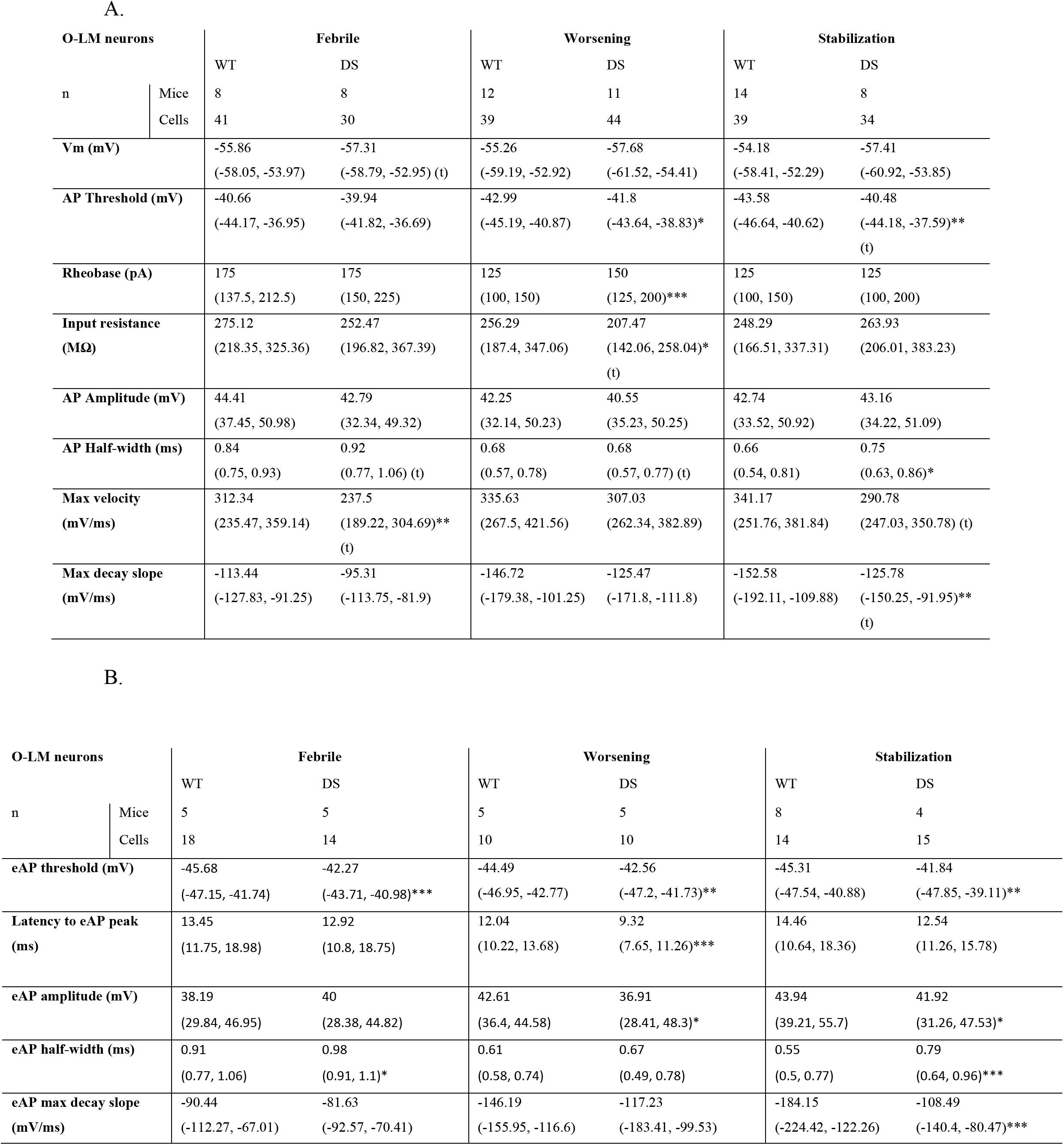
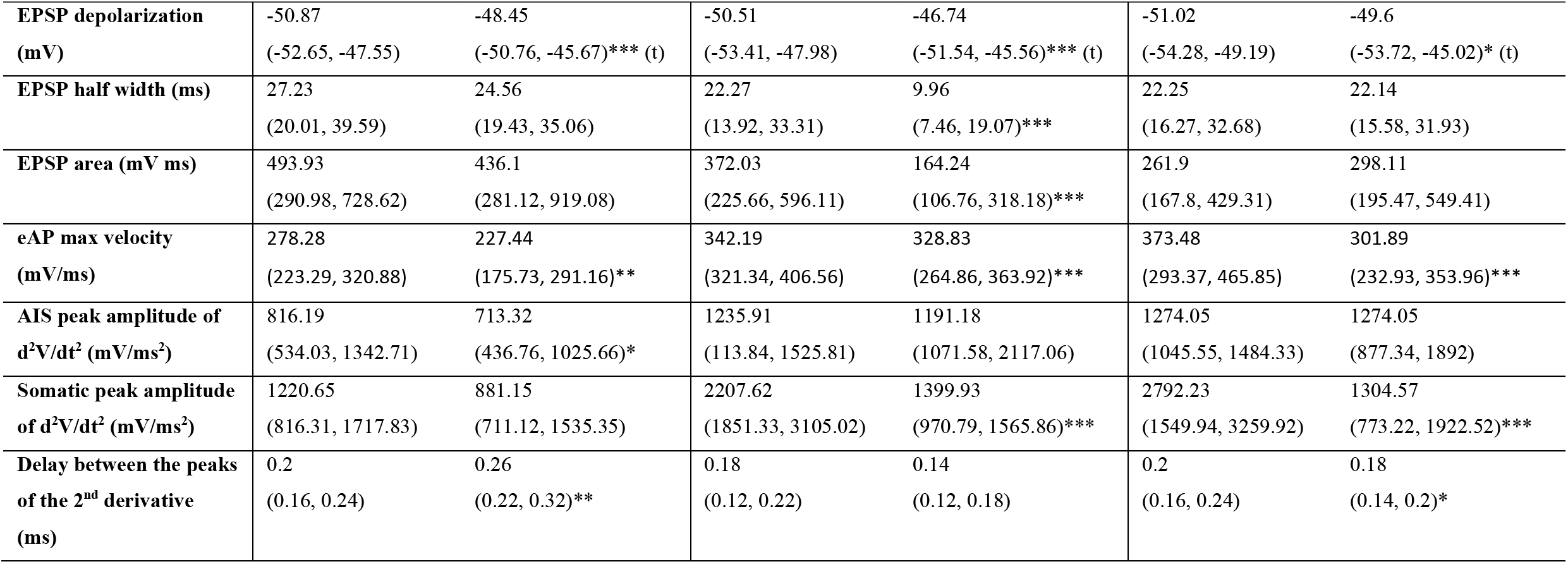
**Electrophysiological properties of O-LM interneurons**, measured in response to direct current injection through the patch pipette (A) or to synaptic stimulation of the Alveus (B). Most of the data did not distribute normally, and are donated as median and range (25%, 75%). Statistical analysis was performed using the Mann - Whitney Rank Sum Test. In cases that the data distributed normally, Student’s t-test was used indicated by (t).

Together, while these data support the involvement of inhibitory neuron dysfunction during the worsening and stabilization stages of Dravet. The febrile state evidenced no signs of reduced interneuron activity.

### Increased excitability of CA1 pyramidal neurons at the febrile stage

Increased activity of excitatory neurons was suggested in dissociated neurons of DS mice (Mistry et al., 2014) and patient-derived neurons (Liu et al., 2013). Nonetheless, no signs of increased firing have yet been found in brain slices (Favero et al., 2018; Rubinstein et al., 2015b; Tai et al., 2014). However, brain slice recordings focused on the worsening and stabilization stages, while the function of excitatory hippocampal neurons during the febrile stage remained untested.

To our surprise, a small increase was evident in the number of APs fired by CA1 pyramidal neurons during the febrile stage (Fig. 4A, B). As opposed to inhibitory neurons, this increase was not detected with low intensity stimulus, but only when the whole curve was included (Fig. 4).

**Figure 4.**
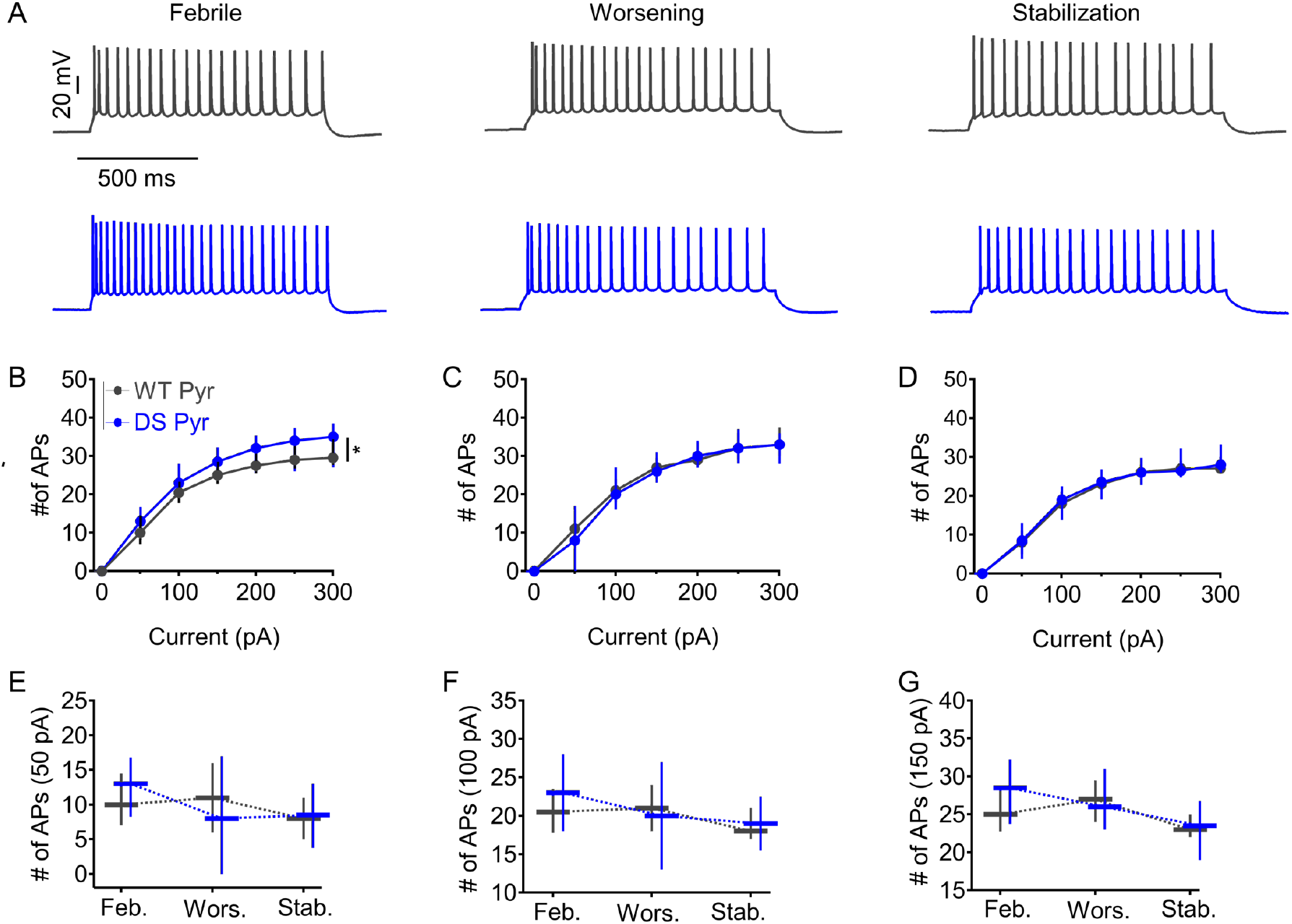
Transient increase in the firing of CA1 pyramidal neurons. (A) Sample traces of whole-cell current clamp recordings from WT (dark gray) and DS (blue) CA1 pyramidal neurons upon injection of 100 pA through the patch pipette. (B-D) AP counts in response to 1 s depolarizing current injection (also corresponding to firing frequency), at the indicated intensities, during the febrile (B), worsening (C) and stabilization (D) stages. Symbols represent the medians; vertical line segments under and above the median represent the 25 and 75 percentile ranges, respectively. Statistical analysis used Two-Way Repeated Measures ANOVA, followed by Holm-Sidak*post-hoc* analysis. (E-G) Number of APs generated in response to 50 pA (E), 100 pA (F) or 150 pA (G), in WT and DS neurons during the three stages of Dravet. Data are presented as median (horizontal midlines) and range (vertical line segments under and above the median signify the 25 and 75 percentile ranges, respectively). WT: n=16-37; DS: n=18-44 (see Table 2A for more details). \**P* < 0.05.

Despite the small increase in firing frequency, the parameters of single APs at rheobase were similar between WT and DS neurons at the febrile stage (Fig. 5 and Table 2A). In contrast, at the worsening stage, the threshold and rheobase were increased, suggesting a mild reduction in the excitability of CA1 pyramidal neurons at the onset of spontaneous seizures (Fig. 5 and Table 2A). Upon the stabilization stage, AP parameters were again similar to those of WT neurons (Figs. 4A, D, 5 and Table 2A).

**Figure 5.**
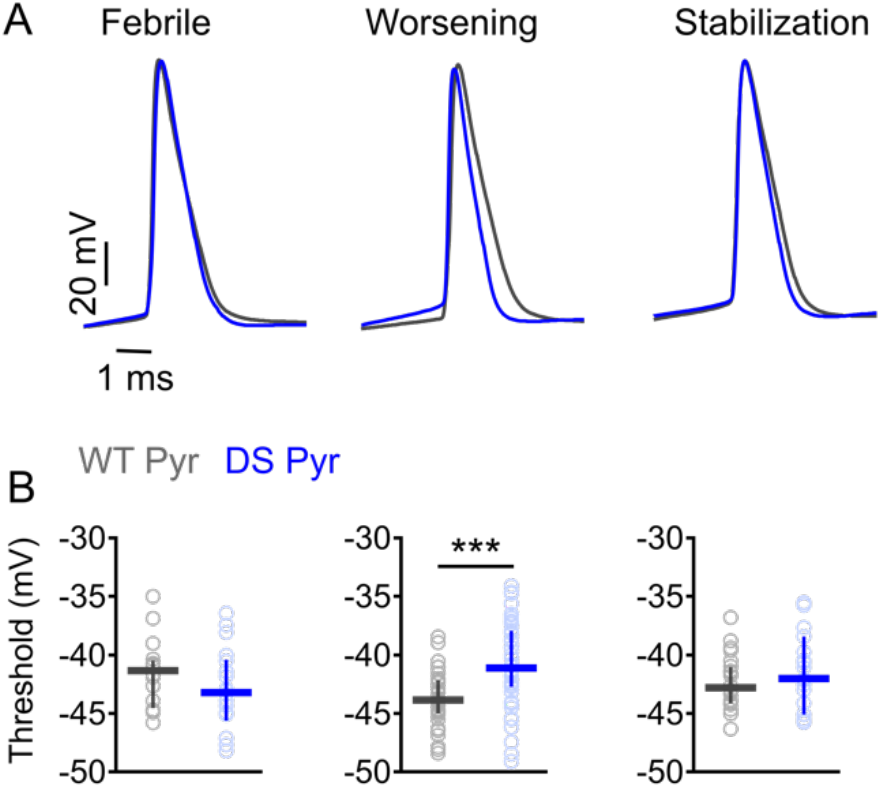
Increased threshold for AP in CA1 pyramidal neurons during the worsening stage. (A) Representative APs from WT (dark gray) and DS (blue) CA1 pyramidal neurons at different ages. Firing was induced by 10 ms depolarizing current injected through the patch pipette at rheobase. (B) AP threshold at the different stages of Dravet as indicated in A. Horizontal midlines represent medians; vertical line segments under and above the median represent 25 and 75 percentile ranges, respectively. Statistical analysis was performed using Mann-Whitney Rank Sum Test. WT: n=16-37; DS: n=18-44 (see Table 2A for more details). \*\*\**P* < 0.001.

**Table 2:**
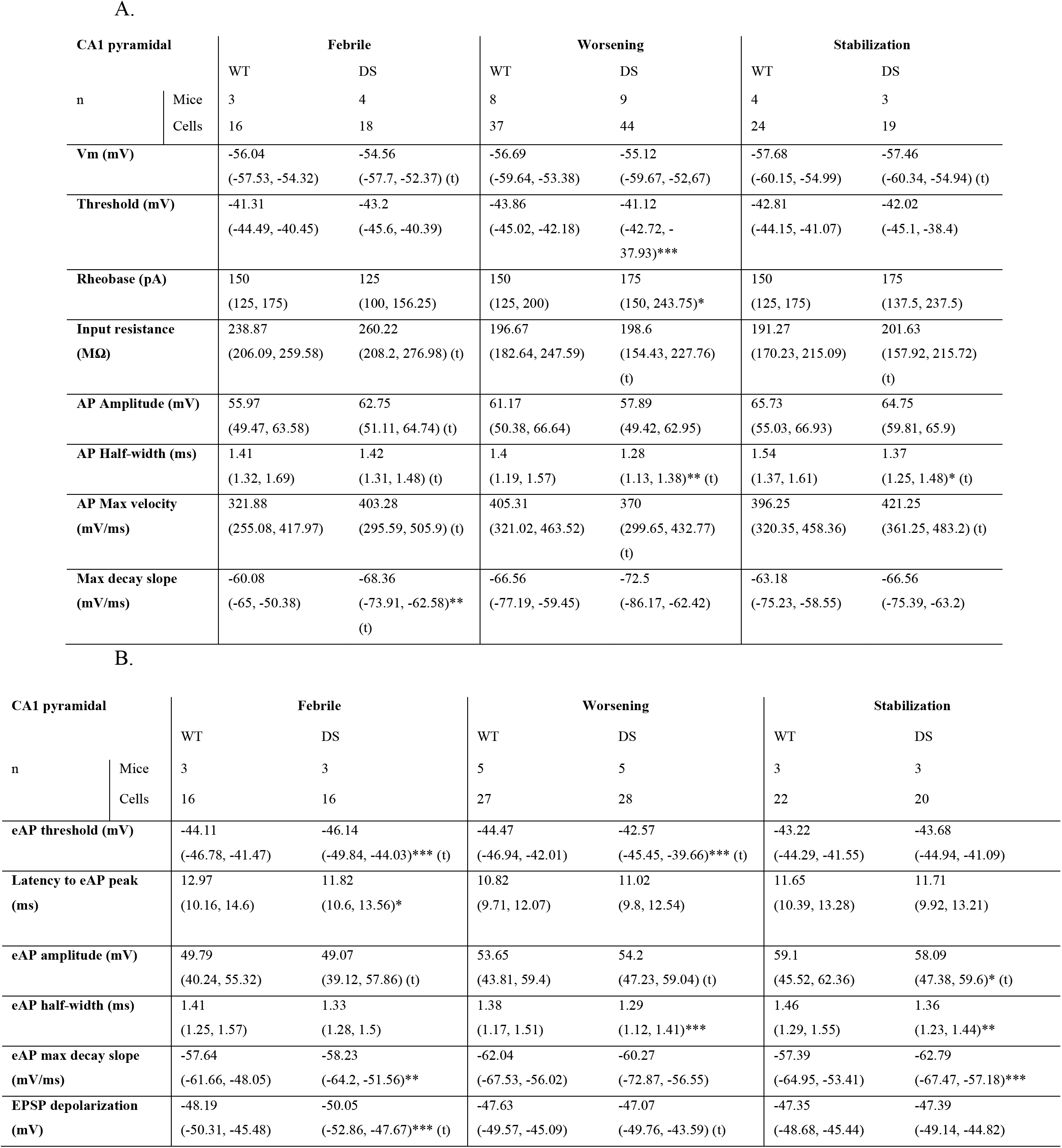
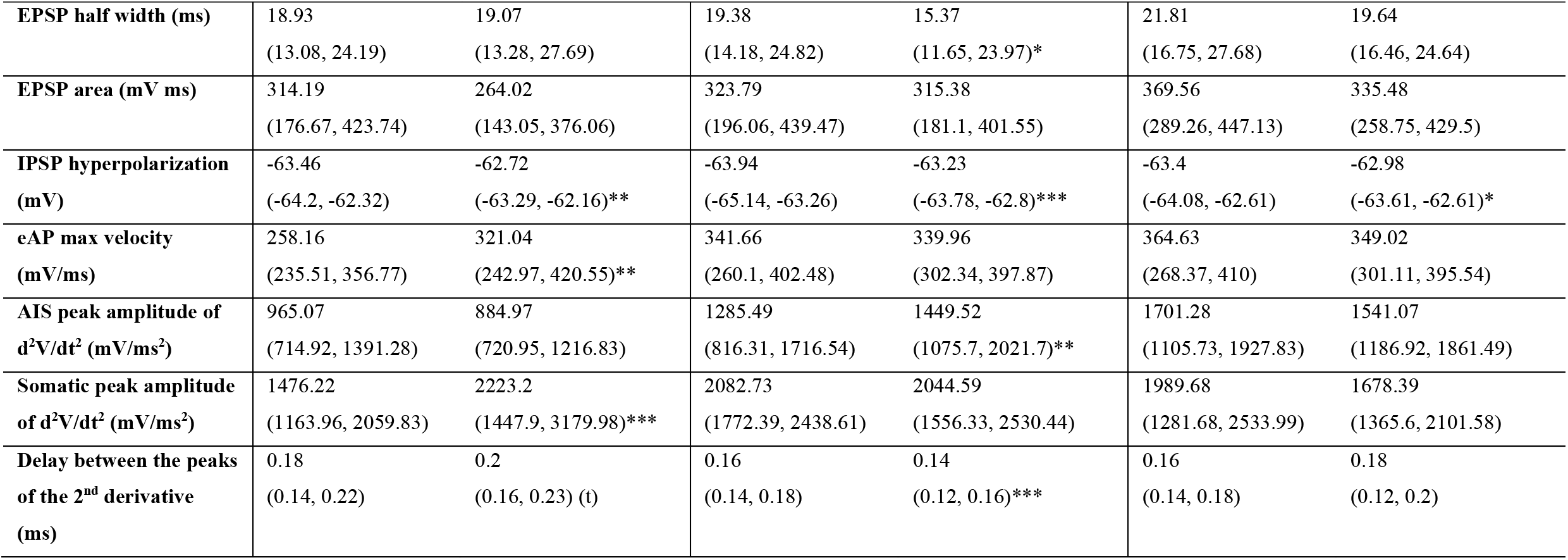
**Electrophysiological properties of CA1 pyramidal neurons** measured in response to direct current injection through the patch pipette (A) or to synaptic stimulation of Shaffer collaterals (B). Most of the data did not distribute normally, and are donated as median and range (25%, 75%). Statistical analysis was performed using the Mann - Whitney Rank Sum Test. In cases that the data distributed normally, Student’s t-test was used indicated by (t).

Together, the firing of CA1 pyramidal neurons is transiently increased during the febrile stage, with correction and signs of reduced excitability at the worsening stage.

### Decreased threshold for synaptically evoked APs in excitatory neurons and increased threshold in interneurons during the febrile stage

The advantage of current injection through the whole cell patch pipette is that the stimulus (the amount of injected current) is fully controlled. Nevertheless, depolarizing inputs are rarely located on the cell soma, and neurons usually operate around rheobase, rather than sustained firing. Moreover, Na_v_1.1 sodium channels are expressed in the cell soma of both excitatory and inhibitory neurons (Ogiwara et al., 2013; Westenbroek et al., 1989), where they contribute towards amplification of synaptic depolarizations (Carter et al., 2012; Magee, 2000; Rubinstein et al., 2015b; Stuart and Sakmann, 1995). Therefore, we reasoned that firing in response to evoked synaptic activity, in which excitatory inputs are received by the dendrites, then integrated and transported through the cell body to the axon initial segment (AIS) (Baranauskas et al., 2013; Magee, 2000; Martina et al., 2000), may highlight additional functional changes between WT and DS.

To normalize the stimulus strength between slices and genotypes, the stimulation was set to produce a firing probability of 50% in the recorded post-synaptic cell, resulting in five synaptically evoked APs (eAPs) and five near-threshold excitatory post-synaptic potentials (EPSPs) out of 10 stimuli at 1 Hz (Rubinstein et al., 2015b). Firing in CA1 pyramidal neurons was evoked by stimulation of the Schaffer collaterals, while antidromic activation of CA1 pyramidal neurons, via stimulation of the Alveus, triggered responses in O-LM neurons (Fig. S1C, D).

During the febrile stage, CA1 pyramidal neurons in DS mice had lower threshold for eAPs (Figs. 6A, B, 9A and Table 2B) and reduced EPSP depolarization (Fig. 6D, E). Moreover, the latency to spike was shorter (Table 2B), together with faster maximal eAP velocity (maximal dV/dt) (Fig. 6C and Table 2B). NaV channels contribute to AP backpropagation. Thus, examination of the second derivative of the AP and quantification of the AIS and somatodendritic portions, can provide an indirect estimate of sodium conduction (Hu et al., 2009; Spratt et al., 2019). Here, examination of the second derivative of the eAPs showed increased acceleration of the late, somatodendritic portion (Fig. 6G-I and Table 2B), suggesting increased sodium conductance in this region.

**Figure 6.**
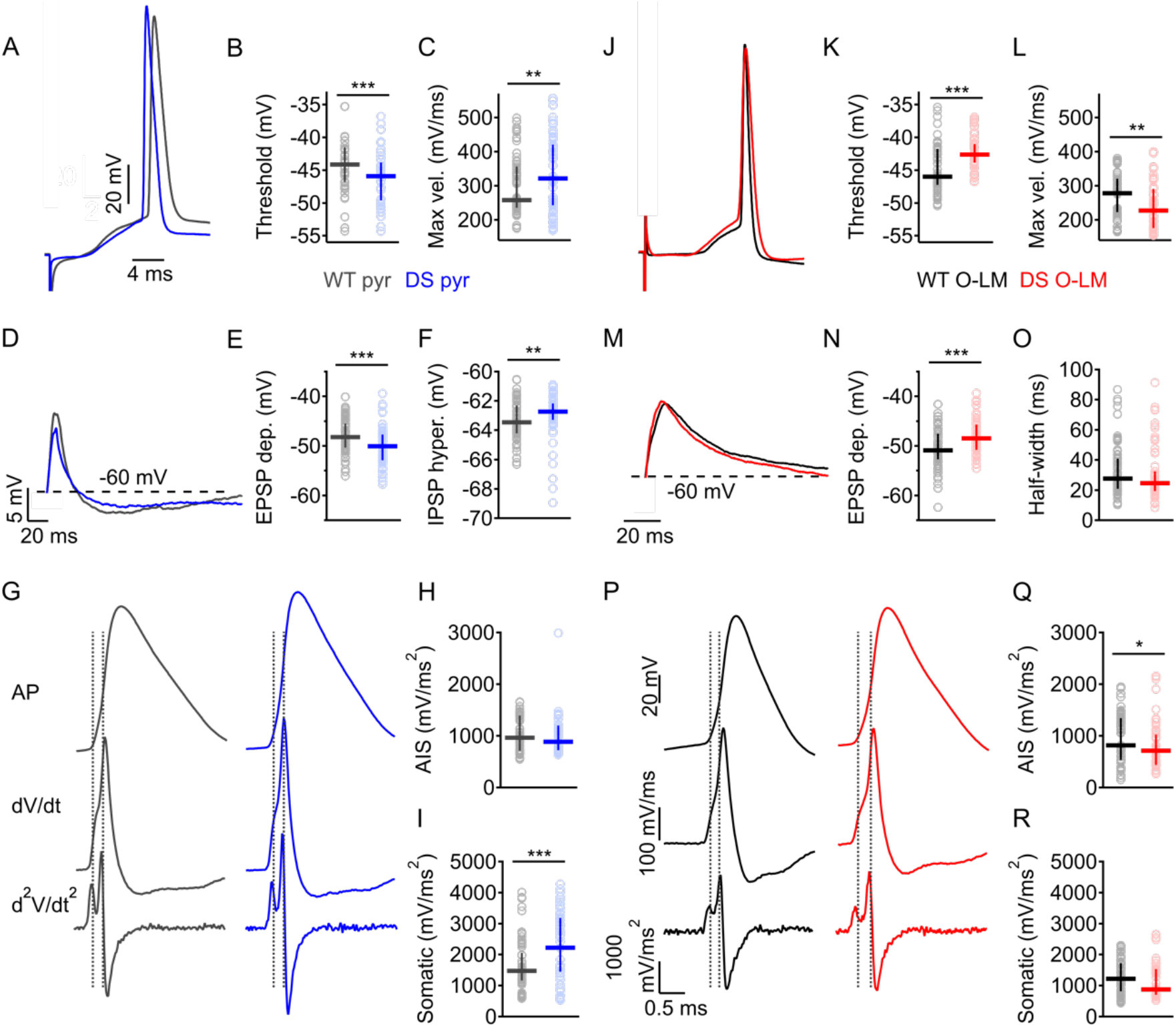
Increased CA1 pyramidal neuron excitability and reduced function of O-LM interneurons during the febrile stage. Synaptically evoked responses (eAPs and subthreshold responses) in CA1 pyramidal neurons were evoked by stimulating the Schaffer collaterals (A-I). eAPs and EPSPs in O-LM interneurons were triggered by stimulating the Alveus (J-R). Stimulation strength was set to produce firing probability of 50% during trains of 10 stimuli at 1 Hz. (A-I) Properties of synaptically evoked responses in WT (dark gray) and DS (blue) CA1 pyramidal neurons during the febrile stage. (A) Representative traces of eAPs at P14. (B) eAP threshold and (C) maximal eAP velocity (maximal rising slope). (D) Representative traces of synaptically evoked subthreshold responses. (E-F) Quantification of the EPSP depolarization (E) and maximal IPSP hyperpolarization (F). (G) eAPs of pyramidal neurons (top panel) and corresponding first (dV/dt; middle panel) and second (d^2^V/dt^2^; bottom panel) derivatives. (H-I) Quantification of the maximal acceleration rates: first AIS acceleration rate (H), and second somatodendritic acceleration rate (I). The horizontal lines represent the medians and the lower and upper lines depict the 25 and 75 percentiles range, respectively. WT pyramidal neurons: n=16; DS pyramidal neurons: n=16. See Table 2B for more information on statistical analysis. (J-R) Properties of synaptically evoked responses in WT (black) and DS (red) O-LM interneurons. (J) Representative traces of eAPs at P14. (K) eAP threshold and (L) maximal eAP velocity. (M) Representative traces of EPSPs. (N) EPSP depolarization and (O) EPSP half-width. (P) Representative eAPs of O-LM interneurons (top panel) and their corresponding first (dV/dt; middle panel) and second (d^2^V/dt^2^; bottom panel) derivatives. (Q-R) Quantification of the maximal acceleration rates: first AIS acceleration rate (Q) and second somatodendritic acceleration rate (R). The horizontal lines represent the medians and the lower and upper lines depict the 25 and 75 percentiles range, respectively. WT O-LM interneurons: n=18; DS O-LM interneurons: n=14. See Table 1B for more information on statistical analysis. *P < 0.05, \*\**P* < 0.01, \*\*\**P* < 0.001.

Schaffer collaterals stimulation also activate PV- and CCK-positive interneurons that mediate feedforward axosomatic inhibition of CA1 pyramidal cells, as well as O-LM cells, which provide dendritic feedback inhibition (Maccaferri and McBain, 1995; Milstein et al., 2015; Scanziani and Pouille, 2004). Quantification of the maximal synaptic hyperpolarization, that follows the EPSP, indicated a small, but significant, reduction in CA1 neurons of DS mice (Figs. 6D, F, 9C and Table 2B). This small reduction may be attributed to impaired activity of inhibitory neurons, alteration in the reversal potential of chloride (Yuan et al., 2019), or both.

We next examined the firing of O-LM interneurons in response to synaptic activation, again setting the strength of the presynaptic stimulation to produce firing probability of 50% (Fig. S1C). The AIS in O-LM neurons is located over 100 μm away from the soma. Therefore in these cells sodium conductance are particularly important for synaptic boosting and fast somatodendritic backpropagation of APs (Martina et al., 2000; Rubinstein et al., 2015a, 2015b). During the febrile stage, O-LM neurons in DS mice had elevated threshold for eAP (Figs. 6J, K, 9H and Table 1B), and accordingly, the EPSP depolarization was also increased (Fig. 6M, N, Table 1B). Analysis of the first and second derivatives of the eAPs revealed reduced maximal eAP velocity (maximal dV/dt) (Figs. 6L, 9K and Table 1B), associated with reduced acceleration of the early, AIS-related phase of the eAPs, while the rate of the late, somatodendritic component, was unchanged (Figs. 6P-R, 9L-M and Table 1B). Moreover, during the febrile stage, the time delay between the two peaks of the second derivative was longer in O-LM neurons of DS mice, indicating a more distal eAP initiation site (Figs. 6P, 9N and Table 1B). In pyramidal neurons, distal AIS positioning, without further change in sodium conductance, was sufficient to cause increased AP threshold and reduced acceleration of the AIS component (Wefelmeyer et al., 2015). Similarly, increased threshold in O-LM cells can be caused by the distal initiation site, reduced NaV conductance, or both. To distinguish between these possibilities, we compared the thresholds in WT and DS neurons that had similar delay between the peaks of the second derivative. Within this subset of data, the threshold for eAP was still elevated, suggesting that reduced sodium conductance, in addition to a more distal eAP initiation, contributes to increased threshold and slower eAP velocity in DS mice (Fig. S2A, B).

Together, during the febrile stage, in which the epileptic phenotypes are mild, there are opposing neuronal changes in both excitatory and inhibitory neurons, with reduced threshold for CA1 pyramidal neurons and increased threshold in O-LM inhibitory neurons.

### Reduced inhibition governs the worsening stage

At the worsening stage, the hyperexcitability of excitatory CA1 neurons was completely overturned. The threshold for eAPs became more depolarized (Figs. 7A, B, 9A and Table 2B), and the duration of the EPSPs was slightly reduced, signifying a narrowing of the time window for synaptic integration (Fig. 7D, E and Table 2B). Additionally, the increased maximal eAP velocity, which was observed during the febrile stage, was corrected (Figs. 7C, 9D. and Table 2B). Analysis of the second derivative indicated a more proximal eAP initiation site in CA1 DS pyramidal neurons, together with increased acceleration of the early, AIS eAP component (Figs. 7G, H, 9E, G and Table 2B). These findings seem in conflict, as elevated threshold is often associated with reduced sodium conductance, while enhanced AIS acceleration rate can indicate increased activity of NaV channels. Moreover, suboptimal AIS positioning can further affect these parameters (Baranauskas et al., 2013). If proximal eAP initiation site is responsible for the increased AIS acceleration, neurons with similar eAP initiation sites (as indicated by similar delays between the two peaks of the second derivative) will have comparable AIS acceleration rates. Indeed, among such cells, the early eAP acceleration rates were similar (Fig. S2C, D). Thus, more proximal eAP initiation site, rather than increased NaV conductance, is probably responsible for the enhanced AIS acceleration (Fig. 7H). Nonetheless, the threshold among these cells was still elevated (Fig S2C, D), indicating that additional alteration of ion channels function may further contribute to the increased threshold.

**Figure 7.**
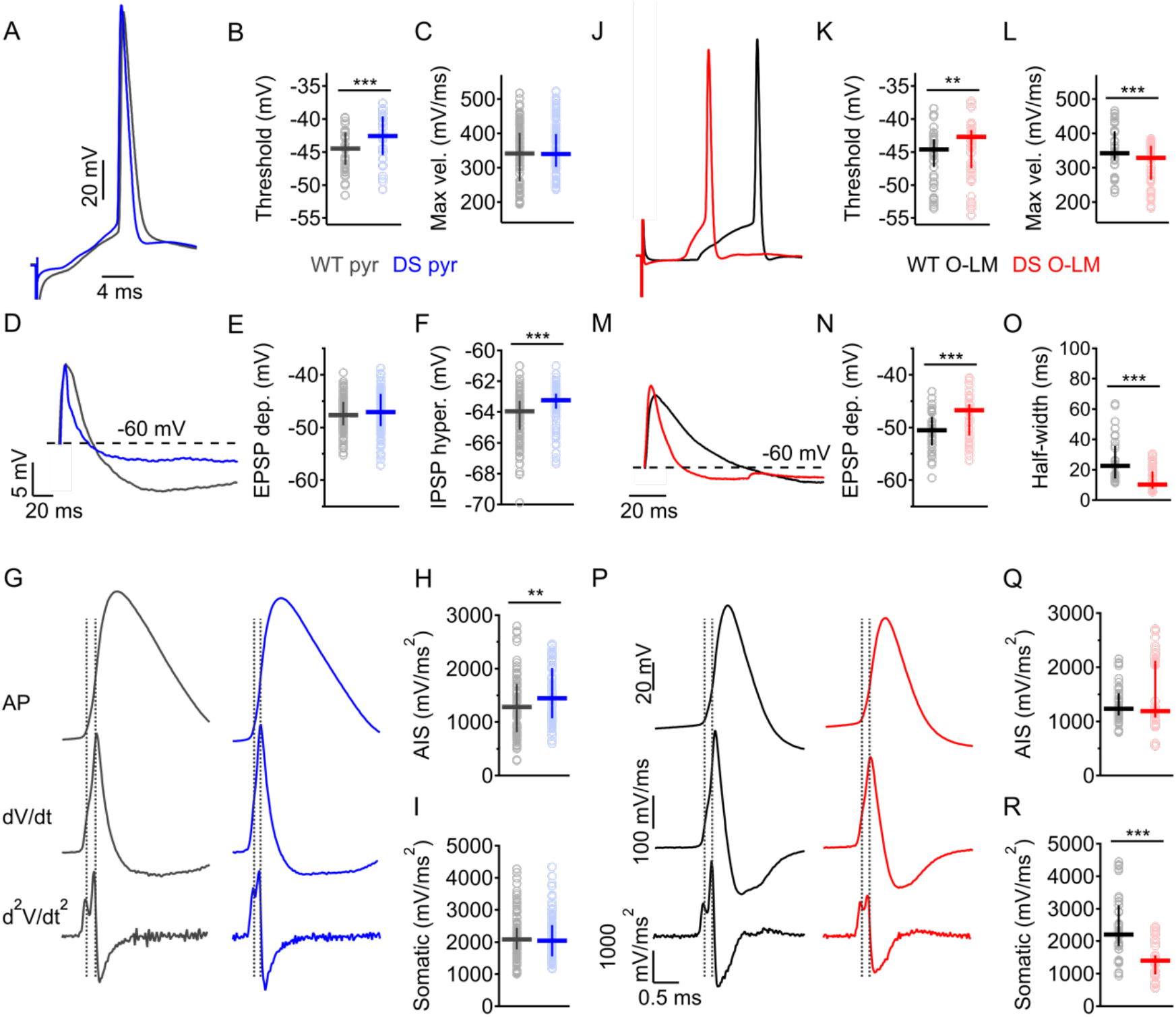
Reduced excitability of CA1 pyramidal neurons and O-LM interneurons during the worsening stage. Synaptically evoked responses (eAPs and subthreshold responses) in CA1 pyramidal neurons were evoked by stimulating the Schaffer collaterals (A-I). eAPs and EPSPs in O-LM interneurons were triggered by stimulating the Alveus (J-R). Stimulation strength was set to produce firing probability of 50% during trains of 10 stimuli at 1 Hz. (A-I) Properties of synaptically evoked responses in WT (dark gray) and DS (blue) CA1 pyramidal neurons during the worsening stage. (A) Representative traces of eAPs at P21. (B) eAP threshold and (C) maximal eAP velocity (maximal rising slope). (D) Representative traces of synaptically evoked subthreshold responses. (E-F) Quantification of the EPSP depolarization (E) and maximal IPSP hyperpolarization (F). (G) eAPs of pyramidal neurons (top panel) and corresponding first (dV/dt; middle panel) and second (d^2^V/dt^2^; bottom panel) derivatives. (H-I) Quantification of the maximal acceleration rates: first AIS acceleration rate (H), and second somatodendritic acceleration rate (I). Horizontal lines represent medians, and vertical line segments under and above the medians depict 25 and 75 percentile ranges, respectively. WT pyramidal neurons: n=27; DS pyramidal neurons: n=28. See Table 2B for more information on statistical analysis. (J-R). Properties of synaptically evoked responses in WT (black) and DS (red) O-LM interneurons. (J) Representative traces of eAPs at P21. (K) eAP threshold and (L) maximal eAP velocity. (M) Representative traces of EPSPs. (N) EPSP depolarization and (O) EPSP half-width. (P) Representative eAPs of O-LM interneurons (top panel) and their corresponding first (dV/dt; middle panel) and second (d^2^V/dt^2^; bottom panel) derivatives. (Q-R) Quantification of the maximal acceleration rates: first AIS acceleration rate (Q) and second somatodendritic acceleration rate (R). Horizontal lines represent medians, and vertical line segments under and above the medians depict 25 and 75 percentile ranges, respectively. WT O-LM neurons: n=10; DS O-LM neurons: n=10. See Table 1B for more information on statistical analysis. \**P* < 0.05, \*\**P* < 0.01, \*\*\**P* < 0.001.

Despite the reduction in CA1 pyramidal neuron activity, inhibition at the worsening stage was severely impaired. The maximal hyperpolarization of CA1 pyramidal neurons was reduced (Fig. 7D, F and Table 2B). Furthermore, direct examination of synaptically evoked firing of O-LM interneurons demonstrated a dramatic reduction in their excitability, with increased threshold for eAP (Fig. 7J, K and Table 1B) and reduced maximal rising slope (Fig. 7L and Table 1B), together with reduced acceleration of the somatodendritic component of the eAP (Fig. 7P, R, and Table 1B). Furthermore, in agreement with a more depolarized eAP threshold, the maximal EPSP depolarization was increased (Figs. 7M, N and Table 1B), while the duration of the EPSP was markedly shorter, dramatically reducing the likelihood for temporal integration (Figs. 7M, O, 9J and Table 1B).

Together, these data indicate that disinhibition governs the onset of spontaneous seizures, with marked reduction in the excitability of O-LM neurons. Notably, however, the activity of CA1 pyramidal neurons at this age is also reduced, possibly ameliorating circuit hyperexcitability.

### Improved function of O-LM inhibitory neurons during the stabilization stage of Dravet

The firing and EPSP properties of CA1 pyramidal neurons in WT and DS mice were similar during the stabilization stage (Figs. 8A-E, 9A, B, D and Table 2B), suggesting complete recovery of pyramidal neuron activity at this age. Nonetheless, the hyperpolarization of the disynaptic inhibition of CA1 excitatory neurons was still reduced (Figs. 8F, 9C and Table 2B).

**Figure 8.**
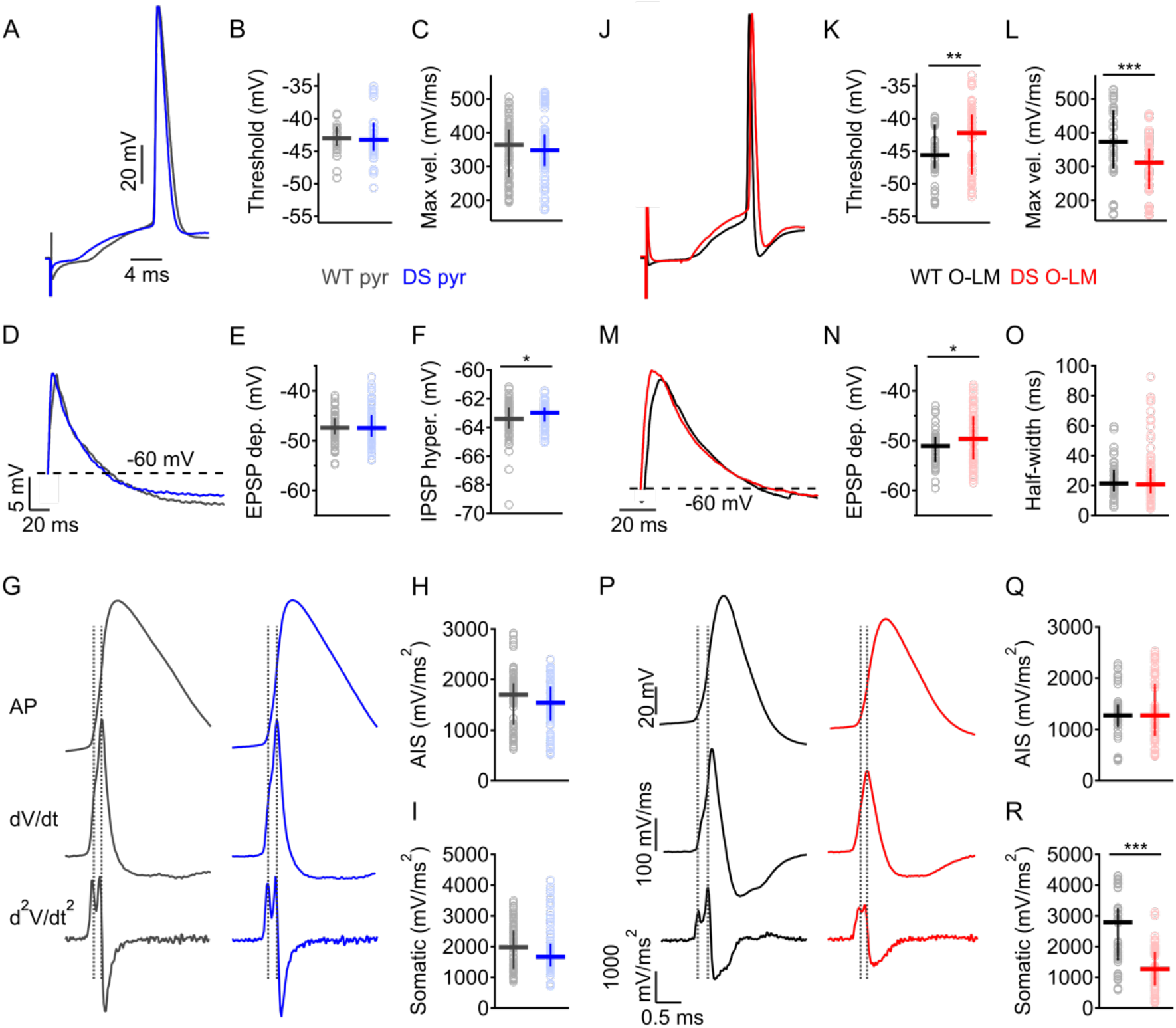
Impaired function of O-LM interneurons during the stabilization stage. Synaptically evoked responses (eAPs and subthreshold responses) in CA1 pyramidal neurons, were evoked by stimulating the Schaffer collaterals (A-I). eAPs and EPSPs in O-LM interneurons were triggered by stimulating the Alveus (J-R). Stimulation strength was set to produce firing probability of 50% during trains of 10 stimuli at 1 Hz. (A-I) Properties of synaptically evoked responses in WT (dark gray) and DS (blue) CA1 pyramidal neurons during the stabilization stage. (A) Representative traces of eAPs at P35. (B) eAP threshold and (C) maximal eAP velocity (maximal rising slope). (D) Representative traces of synaptically evoked subthreshold responses. (E-F) Quantification of the EPSP depolarization (E) and maximal IPSP hyperpolarization (F). (G) eAPs of pyramidal neurons (top panel) and corresponding first (dV/dt; middle panel) and second (d^2^V/dt^2^; bottom panel) derivatives. (H-I) Quantification of the maximal acceleration rates: first AIS acceleration rate (H), and second somatodendritic acceleration rate (I). Horizontal lines represent medians, and vertical line segments under and above the medians depict 25 and 75 percentile ranges, respectively. WT pyramidal neurons: n=22; DS, n=20. See Table 2B for more information on statistical analysis. (J-R). Properties of synaptically evoked responses in WT (black) and DS (red) O-LM interneurons. (J) Representative traces of eAPs at P35. (K) eAP threshold and (L) maximal eAP velocity. (M) Representative traces of EPSPs. (N) EPSP depolarization and (O) EPSP half-width. (P) Representative eAPs of O-LM interneurons (top panel) and their corresponding first (dV/dt; middle panel) and second (d^2^V/dt^2^; bottom panel) derivatives. (Q-R) Quantification of the maximal acceleration rates: first AIS acceleration rate (Q) and second somatodendritic acceleration rate (R). Horizontal lines represent medians, and vertical line segments under and above the medians depict 25 and 75 percentile ranges, respectively. WT O-LM neurons: n=14; DS O-LM neurons: n=15. See Table 1B for more information on statistical analysis. \**P* < 0.05, \*\**P* < 0.01, \*\*\**P* < 0.001.

In contrast, examination of the properties of O-LM neurons during the stabilization stage revealed only a partial correction of their activity. While increased eAP threshold (Figs. 8J, K, 9H and Table 1B), increased EPSP depolarization (Fig. 8M, N and Table 1B) and reduced eAP velocity (Figs. 8L, 9K and Table 1B) persisted, the duration of the EPSPs was corrected, resulting in an overall depolarization time similar to that of WT neurons (Figs. 8M, O, 9J and Table 1B). Additionally, the duration of the eAP was increased, together with a slower decay (Table 1B), which may contribute to increased GABA release (Bean, 2007). Analysis of the second derivative of the O-LM cells eAP showed shorter delay between the first and second peaks, indicating a slightly more proximal eAP initiation site (Figs. 8P, 9N). Nevertheless, the acceleration rate of the somatodendritic component was still reduced, without much change from the worsening stage (Figs. 8P, R, 9M and Table 1B).

Together, our results show that while CA1 pyramidal neurons regain normal properties, reduced function of inhibitory neurons persists through the stabilization stage, although with some improvement relative to the worsening stage.

**Figure 9.**
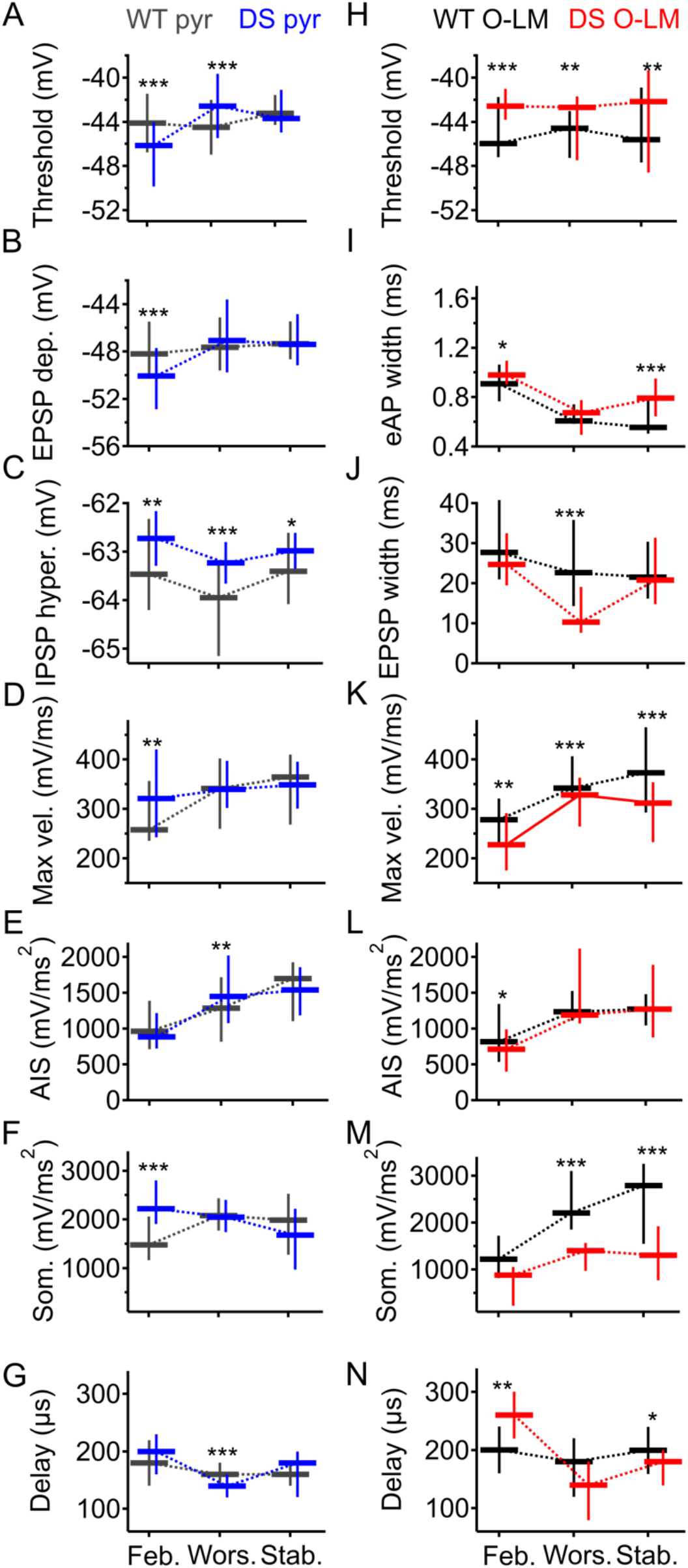
Time course of changes in the properties of CA1 pyramidal and O-LM neurons. (A-G) Developmental changes in the properties of synaptically evoked responses in WT (gray) and DS (blue) CA1 pyramidal neurons. (A) eAP threshold, (B) EPSP depolarization, (C) IPSP hyperpolarization, (D) maximal eAP velocity, (E) AIS acceleration rate, (F) somatodendritic acceleration rate and (G) the delay between the two peaks of the eAP second derivative. WT pyramidal neurons: n=16-27; DS pyramidal neurons: n=16-28 (see Table 2B for more details). (H-N) Developmental changes in the properties of synaptically evoked responses in WT (black) and DS (red) O-LM interneurons. (H) eAP threshold, (I) eAP half-width, (J) EPSP half-width, (K) eAP maximal velocity, (L) AIS acceleration rate, (M) somatodendritic acceleration rate and (N) the delay between the two peaks of the eAP second derivative. Statistical analysis utilized Two-Way ANOVA, followed by Holm-Sidak *post-hoc* analysis (See Table S1 for more information on statistical comparison within genotypes during development). Data are presented as median (horizontal midlines) and range (vertical line segments under and above the median signify the 25 and 75 percentile ranges, respectively). WT O-LM interneurons: n=10-18; DS O-LM interneurons: n=10-15. (see Table 1B for more details). *P* < 0.05, \*\**P* < 0.01, \*\*\**P* <0.001.

## Discussion

Dravet epilepsy manifests with a dynamic time course, beginning with susceptibility to febrile seizures, progression to spontaneous seizures and profound premature death, which is followed by some amelioration of the epileptic phenotypes. Here we provide a comprehensive analysis of the activity of CA1 pyramidal neurons and O-LM interneurons at each of these three different stages, demonstrating evolving mutual changes in their activity.

### Susceptibility to thermally induced seizures correlates with the risk of premature death

With disease progression from the febrile stage through the worsening to the stabilization stage, DS mice harboring the missense *Scn1a^A1783V^* mutation, recapitulate the epileptic phenotypes of Dravet. Importantly, DS mice exhibit neither spontaneous seizures nor premature mortality during their first weeks of life, yet progress to high incidences of seizure activity and mortality during the fourth week (~P20-P28) (Fig. 1A). Notably, the onset of premature death and overall mortality in these *Scn1a^A1783V^* mice is very similar to that of other DS mice on the C57BL/6J genetic background (Kang et al., 2018; Ogiwara et al., 2007; Styr et al., 2019; Tsai et al., 2015; Yu et al., 2006). Moreover, as with Dravet patients, susceptibility to thermally induced seizures precedes the onset of spontaneous seizures (Dravet, 2011; Oguni et al., 2001).

Though spontaneous seizures and premature death rarely occurred prior to P18 (Fig. 1A) (Mistry et al., 2014; Ogiwara et al., 2007; Styr et al., 2019; Yu et al., 2006), the high sensitivity to thermal induction of seizures, with all of the seizures occurring below 39.5 °C (Fig. 1B, C), indicates that neuronal changes have already commenced during this mostly asymptomatic febrile stage. Of note, susceptibility to early febrile seizures (at P14-P16) was demonstrated before (Hawkins et al., 2017). However, unlike the DS mice that were used here, Hawkins et al. used mice on the mixed C57BL/6J:129S6/SvEvTac genetic background, which had thermally induced seizures at high temperatures (average temperature of 42 °C) (Hawkins et al., 2017). Thus, in addition to modifying the survival of DS mice (Kang et al., 2018; Yu et al., 2006), genetic background also alters the febrile stage of Dravet.

During the course of Dravet, the temperature of seizure induction changed in correlation with the risk of premature death (Fig. 1B). During the febrile and stabilization stages, the occurrence of premature death was lower and seizure temperature was relatively high. However, at the worsening stage, which is characterized by profound mortality, thermal induction of seizures was initiated by a small elevation in body core temperature (Fig. 1B, C).

Together, the commercially available DS mice, harboring the missense *Scn1a^A1783V^* mutation, maintained on the pure C57BL/6J genetic background fully recapitulate the three stages of Dravet disease. These mice present with the febrile, worsening and stabilization stages similarly to DS mouse models that are based on *Scn1a* truncation mutations.

### Increased excitability of CA1 pyramidal neurons is limited to the febrile stage of Dravet

The involvement of excitatory neurons in Dravet is debatable. On one hand, electrophysiological recordings from brain slices, performed mostly during the worsening stage, did not detect any alterations in their function (De Stasi et al., 2016; Favero et al., 2018; Rubinstein et al., 2015b). In contrast, somatic sodium currents, in both dissociated hippocampal pyramidal cells (Mistry et al., 2014) and Dravet patient-derived neurons (Liu et al., 2013), were shown to be increased. Moreover, immunocytochemical experiments clearly showed the expression of Na_v_1.1 in excitatory neurons (Ogiwara et al., 2013; Westenbroek et al., 1989). Finally, deletion of *Scn1a* in both excitatory and inhibitory neurons improved the survival compared to mice with deletion only in inhibitory neurons (Ogiwara et al., 2013), indicating that *Scn1a* mutations affect Dravet outcome by perturbing the function of both neuron types.

Here, we examined the activity of CA1 pyramidal neurons at the three stages of Dravet, comparing their firing in response to current injection through the whole cell patch pipette, as well as in response to physiological synaptic stimulation via activation of the Schaffer collaterals. The advantage of the first experimental paradigm is that the amount of injected current is fully controlled, providing known input stimuli. Conversely, synaptic activation, using scaled stimuli, mimics physiological synaptic dendritic depolarization and can further highlight the contribution of Na_v_ channels to active dendrites properties.

Our recordings show a transient increase in the activity of CA1 pyramidal neurons that is limited to the febrile stage. Pyramidal neurons hyperexcitability is indicated by a small increase in the number of APs fired in response to current injection (Fig. 4A, B) and lower threshold for eAPs (Fig. 6A, B and Table 2B). Further analysis of the eAP properties indicated that this enhanced excitability may be related to increased sodium conductance, as suggested by increased maximal eAP velocity (Figs. 6C 9D and Table 2B) and enhanced acceleration of the somatodendritic portion of the eAP (Figs. 6G, I, 9F and Table 2B). The cellular mechanisms contributing to such enhanced function, despite loss of function of Na_v_1.1 channels, are unclear. However, in Dravet patient-derived neurons, as well as in DS mouse dissociated hippocampal pyramidal neurons, increased sodium conductance was attributed to post-translational modifications (Liu et al., 2013; Mistry et al., 2014).

While comparison between firing properties in response to current injection to the soma and synaptic stimulation (Figs. 4, 6A-I and Table 2) show similar trends, the results are more profound using synaptic stimulation. This may be due to the crucial role of Na_v_ channels in active dendritic properties, contributing to boosting or filtering of synaptic depolarization (Carter et al., 2012; Ferrarese et al., 2018; Häusser, 2001; Stuart and Sakmann, 1995). In contrast, when the large soma is directly depolarized, the effect of somatodendritic conductances may be partially bypassed. Nevertheless, during the worsening stage, at the onset of spontaneous seizures, the hyperexcitability of pyramidal neurons is completely overturned, with further indications of reduced excitability. At this age, eAPs showed depolarized threshold, the duration of EPSPs was reduced and the maximal rising slope was similar to that of WT (Figs. 7A-C, 9A, D and Table 2B). We propose that homeostatic changes, prompted by the hyperexcitability of the febrile stage, lead to compensatory reduction in the activity of CA1 pyramidal neurons. These changes likely have a protective role in Dravet, facilitating the reduction of circuit excitation when inhibition breaks down at the onset of spontaneous seizures (see below). Indeed, genetic studies further corroborated the potential protective role of *Scn1a* mutations in excitatory neurons. In DS mice with a *Scn1a* truncation mutation, selective deletion in inhibitory neurons resulted in complete mortality, while deletion in both excitatory and inhibitory neurons reduced the mortality level by 50% (Ogiwara et al., 2013). Similarly, DS mice that harbor the *Scn1a^A1783V^* missense mutation solely within inhibitory neurons do not survive beyond P23 (Kuo et al., 2019), while those with global expression of the mutation, as done here, exhibit a mortality rate of ~50% (Fig. 1A).

### The level of O-LM interneurons dysfunction correlates with the severity of epilepsy

The dysfunction of inhibitory neurons has a major contribution to the etiology and progression of Dravet (Cheah et al., 2012; Favero et al., 2018; Han et al., 2012; Kalume et al., 2013; Ogiwara et al., 2013, 2007; Rubinstein et al., 2015a, 2015b; Tai et al., 2014; Tsai et al., 2015; Yu et al., 2006). Here, we demonstrate that the impaired function of inhibitory O-LM neurons, beginning from the febrile stage through the worsening and stabilization stages, correlates between the level of hampered function and the severity of epilepsy (Figs. 1, 6J-R, 7J-R, 8J-R, 9H-N and Table 1). Deficits that persisted throughout the different stages of Dravet included elevated threshold for eAPs and reduced eAP velocity (Figs. 6J-L, 7J-L, 8J-L, 9H, K and Table 1B). These alterations indicate a mechanism consisting of reduced sodium conductance at all of these disease stages. Despite that, the functional impairment of inhibitory neurons is mostly evident at the onset of seizures. At the worsening stage, in addition to the elevated threshold and reduced eAP velocity, the firing is reduced (Fig. 2), EPSPs are markedly shorter (Figs. 7M, O, 9J and Table 1B), narrowing temporal integration window, and the input resistance is lower (Fig. 3C, D and Table 1A). Firing of PV-positive inhibitory neurons in DS mice was recently shown to depend on the ratio between sodium and potassium currents (Goff and Goldberg, 2019). Similarly, impaired inhibitory function at the worsening stage may be due to a combination of reduced sodium conductance together with developmental upregulation of potassium channels (Downen et al., 1999; Falk et al., 2003; Goldberg et al., 2011). Indeed, the altered input resistance at the worsening stage may indicate enhanced conductance of potassium channels. Moreover, the improved function during the stabilization stage, demonstrated by longer EPSPs and eAPs (Figs. 8M, O, 9J and Table 1B), together with slower eAP decay (Table 1B), is consistent with reduced potassium conductance, which may be a compensatory change aimed to rectify the imbalance between sodium and potassium currents.

## Conclusion

Here we provide a comprehensive analysis of the activity of hippocampal excitatory and inhibitory neurons during the febrile, worsening and stabilization stages of Dravet. Our data indicate increased excitability of pyramidal neurons, that is limited to the febrile stage, and dysfunction of inhibitory neurons at multiple stages of the disease, with correlation between the level of impairment and the severity of epilepsy.

## Material and Methods

### Mice

All animal experiments were approved by the Animal Care and Use Committee (IACUC) of Tel Aviv University. Mice used in this study were housed in a standard animal facility at the Goldschleger Eye Institute at a constant (22°C) temperature, on 12-hour light/ dark cycle, with ad libitum access to food and water.

DS mice harboring the global *Scn1a^A1783V^* mutation were generated by crossing males bearing the conditional *Scn1a^A1783V^* floxed allele (B6(Cg)-*Scn1a*^*tm1.1Dsf*^/J; strain 026133; The Jackson Laboratory) (Kuo et al., 2019; Styr et al., 2019) with CMV-Cre females (B6.C-Tg(CMV-cre)1Cgn/J; strain 006054; The Jackson Laboratory). This breeding scheme was used due to the location of Cre on the X chromosome (Schwenk et al., 1995). These lines were maintained on the pure C57BL/6J genetic background.

The F1 offspring of this cross are WT and DS mice, the latter expressing a global heterozygous *Scn1a^A1783V^* mutation. Both male and female offspring were used. Mice carrying the floxed *Scn1a^A1783V^* allele but lacking Cre expression were not used due to the possibility of spontaneous recombination or flox leakiness.

### Thermal induction of seizures

Thermal induction of seizures was performed in DS mice of different ages: P14-P16 (the febrile stage), P20-P25 (the worsening stage) or P33-P45 (the stabilization stage), as previously described (Oakley et al., 2013, 2009; Rubinstein et al., 2015b). Briefly, the baseline body core temperature of mice was recorded for at least 10 min, allowing the animals to habituate to the recording chamber and rectal probe. Body temperature was then monitored (TCAT-2DF, Physitemp Instruments Inc., Clifton, New Jersey, USA) and increased by 0.5°C every 2 min with a heat lamp, until a generalized tonic-clonic seizure was provoked; the temperature was not increased above 41°C. Seizure temperature was extracted from the video recording. In this test, all of the tested DS mice had generalized tonic-clonic seizures. In contrast, seizure activity could not be induced in any of the WT mice that were heated to 41°C.

### Brain slice electrophysiology

Hippocampal slices were prepared from WT or DS littermates aged P14–P16 (the febrile stage), P21–P24 (the worsening stage) or P31–P35 (the stabilization stage) using standard procedures as before (Rubinstein et al., 2015b). Briefly, mice were deeply anesthetized with isoflurane and decapitated. The brain was then quickly removed into cold slicing solution containing 75 mM sucrose, 87 mM NaCl, 25 mM NaHCO3, 25 mM D-glucose, 2.5 mM KCl, 1.25 mM NaH_2_PO_4_, 0.5 mM CaCl_2_ and 7.0 mM MgCl_2_.

Slicing was performed using a Leica VT1200S vibratome (Leica Biosystems, Wetzlar, Germany). For recording O-LM inhibitory cells, longitudinal slices were used to better preserve the synaptic connections. The brain was cut midline and the hemispheres were glued on their callosal side onto a slope of 45°, forming longitudinal sagittal slices (400 μm thick) (Rubinstein et al., 2015b; Tort et al., 2007). For recording CA1 pyramidal cells, the brain was glued on its ventral side, without any tilt, and horizontal slices were taken (400 μm thick). After sectioning, slices were transferred to a storage chamber with fresh ACSF (containing 125 mM NaCl, 3 mM KCl, 2.0 mM MgCl2, 2.0 mM CaCl2, 1.25 mM NaH2PO4, 26 mM NaHCO3, and 10 mM D-glucose) and incubated for 45 min at 37°C, followed by a 30 min recovery at room temperature. All solutions were saturated with 95% O2 and 5% CO2. The cells were visualized under oblique illumination with near-infrared LED and an upright microscope (SOM; Sutter Instrument, Novato, CA, USA). Stratum oriens O-LM interneurons were identified by their horizontal soma and their characteristic sag (Maccaferri and McBain, 1995; Rubinstein et al., 2015b; Zemankovics et al., 2010); CA1 pyramidal neurons were identified by the alignment of their triangular soma in the thin stratum pyramidale layer (Rubinstein et al., 2015b).

Recordings were obtained at room temperature, using a Multiclamp 700B amplifier (Molecular Devices, San Jose, CA, USA) and Clampex 10.7 software (Molecular Devices). The patch pipette was filled with internal solution containing: 145 mM K-Gluconate, 2 mM MgCl2, 0.5 mM EGTA, 2 mM ATP-Tris, 0.2 mM Na2-GTP and 10 mM HEPES, pH 7.2. When filled with intracellular solution, patch electrode resistance ranged from 4 to 7 MΩ. Access resistance was monitored continuously for each cell. Only cells with access resistance less than 25 MΩ were recorded. The resting membrane potential was set to −60 mV by injection of no more than 50 pA, ranging from −50 to +50 pA. Bridge balance and pipette capacitance neutralization, based on the amplifier circuit, were applied to all current clamp recordings. Data from electrophysiology experiments were analyzed using Clampfit 10.7 (Molecular Devices) and Igor Pro (WaveMetrics, Lake Oswego, OR, USA). Action potential (AP) threshold was defined as the voltage at which the first derivative of the AP waveform (dV/dt) reaches 10 mV/ms. AP amplitude was defined as the overshoot (AP peak relative to 0 mV; Bean, 2007). AP half-width was defined as the time duration of the AP at half of its maximal amplitude. Maximal rising slope (dV/dt) was defined as the maximum value of the first derivative of the AP waveform, at which the velocity of the AP rising phase is at its highest, while maximal decay slope was defined as the lowest, most negative value of the first derivative, at which the velocity of the AP falling phase is highest. The rheobase was determined by injection of incrementing depolarizing current (Δ25 pA) through the patch pipette. Input resistance was averaged from 1 sec long, −50 pA and −100 pA hyperpolarizing current injections through the patch pipette, and measured from the end of the step (following the sag, at steady state).

For synaptically-evoked activity of O-LM interneurons, the stimulating electrode was placed on the alveus, while synaptically-evoked activity of pyramidal neurons was measured by placing the stimulating electrode on the Schaffer collaterals. Stimulation strength was set to produce a firing probability of 50% (i.e., 5 APs and 5 near-threshold excitatory post-synaptic potentials (EPSPs) out of 10 stimuli at 1 Hz), as performed previously (Rubinstein et al., 2015b). Cells which produced 3 to 7 Aps were used for analysis. The EPSP amplitude depolarization was measured as the maximal subthreshold depolarization, while the inhibitory post-synaptic potential (IPSP) was measured as the minimal voltage that followed the EPSP. The EPSP half-width was defined as the duration at half of the maximal amplitude.

### Experimental design and statistical analysis

Statistical analyses were performed using SigmaPlot 12.5 software (Systat Software, San Jose, CA, USA). Most data were not normally distributed, thus are presented as medians and range (25% and 75%). For comparison between WT and DS mice of the same age group, we used the non-parametric Mann-Whitney test (for data that did not distribute normally) or Student’s t-test (for data with normal distribution), as indicated in Table 1 and Table 2. Statistical comparison of the firing in Fig. 2B-D and Fig. 4B-D was conducted using Two-Way Repeated Measures ANOVA followed by Holm-Sidak *post-hoc* analysis. To determine statistical change over time for a specific genotype, we used Two-Way ANOVA followed by Holm-Sidak *post-hoc* analysis. We considered *P* < 0.05 as having reached statistical significance.

## Acknowledgements

We thank I. Fleidervish and B. Attali and Y. Hatin for helpful discussions and critical reading of the manuscript. This study was supported by the Israel Science Foundation grant 1454/17 (MR), Fritz Thyssen Stiftung (10.17.1.023MN) and The Aya Baharav fund, Sackler Faculty of Medicine, Tel Aviv University

