## Supplementary material for "Early hippocampal hyperexcitability followed by disinhibition in a mouse model of Dravet syndrome": Fig. S1, Fig. S2, Table S1

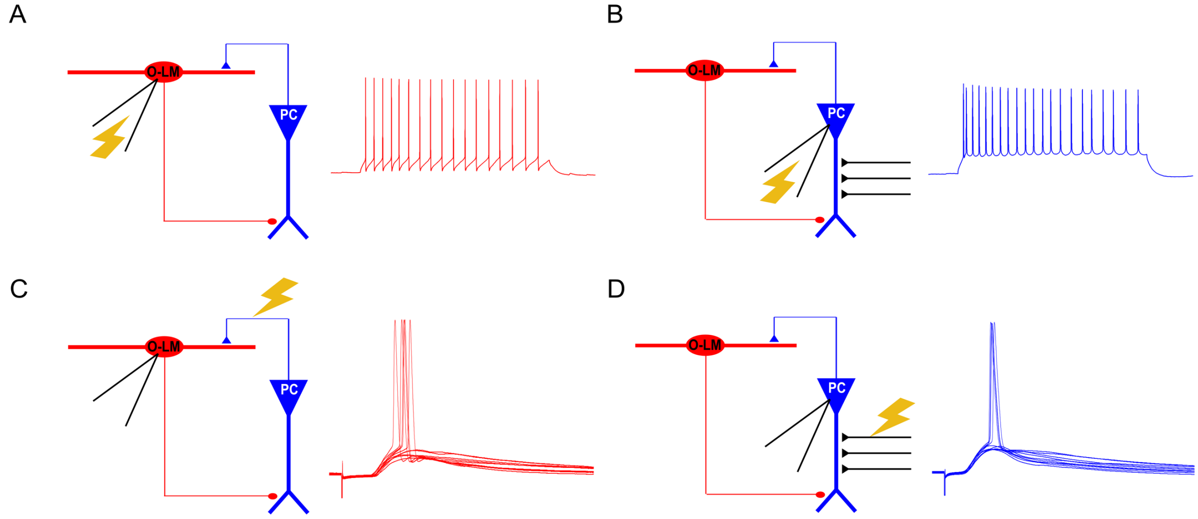


**Fig. S1**. **Schematic drawings of the different forms of recordings performed in the CA1 microcircuit** Pyramidal cells (PC) are illustrated in blue while O-LM inhibitory interneuron are depicted in red. CA1 pyramidal neurons are innervated by the Schaffer collaterals, illustrated in black. O-LM neurons are activated by the axons of CA1 pyramidal neurons (blue). (A-B) Neuronal firing was evoked by current injection through the patch pipette into (A) O-LM interneurons and (B) CA1 pyramidal neurons. (C-D) Neuronal activity was evoked by stimulation of the Alveus (C) or the Schaffer collaterals (D). The stimulation strength was set to produce a firing probability of 50% in the recorded post-synaptic cell, resulting in five synaptically evoked APs (eAPs) and five near-threshold excitatory post synaptic potentials (EPSPs) out of 10 stimuli at 1 Hz.

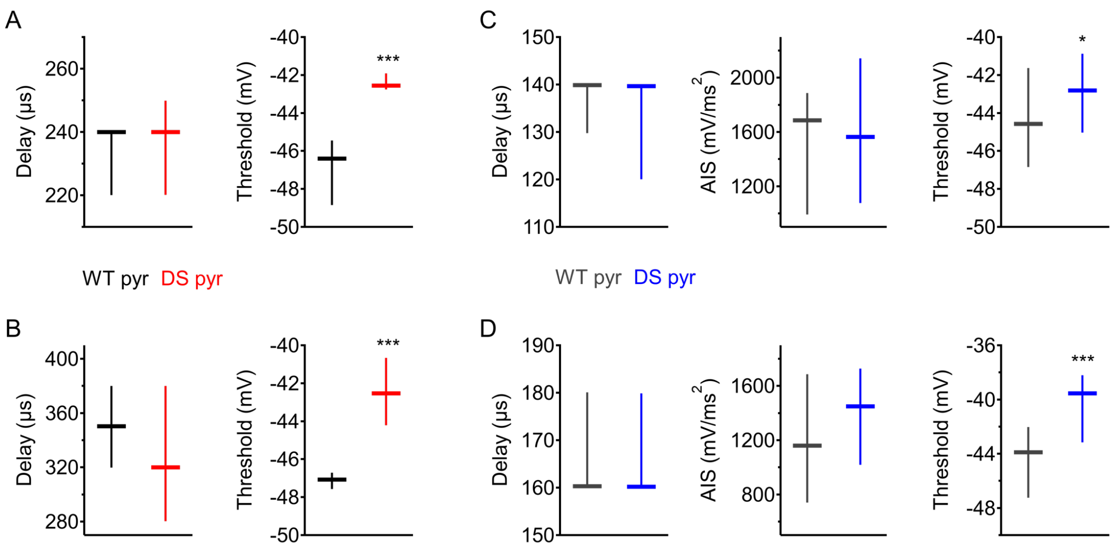


**Fig. S2. Elevated threshold in neurons with similar delay between the first and second peaks of the second derivative**

(A-B) The elevated threshold in O-LM neurons, during the febrile stage, is not related to a more distal AP initiation site (Fig. 9N and Table 1B). Time delay between the AIS and the somatodendritic components of the eAP second derivative (left panel) and eAP threshold (right panel) of WT and DS mice at the febrile stage. Comparison between neurons with similar delay, 220-260 µs (A) and 280-420 µs (B) demonstrated that the elevated threshold is independent of the AP initiation site. (C-D) At the worsening stage, DS CA1 pyramidal neurons had a shorter delay between the two components of the second derivative (Fig. 9G and Table 2B). In addition, the contribution of the first component of the eAP propagation (AIS), was more prominent (Figs. 7H, 9E and Table 2B). Comparison between neurons with similar delay, 120-150 µs (C) and 150-190 µs (D), showed similar AIS acceleration rates as well. Nonetheless, within these neurons, the AP threshold was still elevated. Thus, increased eAP threshold in DS CA1 pyramidal neurons, during the worsening stage, is probably not related to a more proximal AP initiation site. Statistical analysis was performed using Mann-Whitney Rank Sum Test. The horizontal lines represent the medians and the lower and upper lines represent the 25 and 75 percentiles range, respectively. *$P < 0.05$, ***$P < 0.001$.

| | | Interneurons | | | | Pyramidal neurons | | | |
|---|---|---|---|---|---|---|---|---|---|
| | | WT | | DS | | WT | | DS | |
| | Age | P-value | Significance | P-value | Significance | P-value | Significance | P-value | Significance |
| **AP threshold (mV)** | P14 vs. P21 | 0.519 | No | 0.021 | Yes | 0.387 | No | <0.001 | Yes |
| | P21 vs. P35 | 0.938 | No | 0.228 | No | 0.001 | Yes | 0.247 | No |
| | P35 vs. P14 | 0.585 | No | 0.159 | No | 0.046 | Yes | <0.001 | Yes |
| **EPSP Depolarization (mV)** | P14 vs. P21 | 0.900 | No | 0.918 | No | 0.618 | No | <0.001 | Yes |
| | P21 vs. P35 | 0.703 | No | 0.180 | No | 0.660 | No | 0.823 | No |
| | P35 vs. P14 | 0.678 | No | 0.247 | No | 0.526 | No | <0.001 | Yes |
| **IPSP hyperpolarization (mV)** | P14 vs. P21 | | | | | <0.001 | Yes | 0.014 | Yes |
| | P21 vs. P35 | | | | | <0.001 | Yes | 0.125 | No |
| | P35 vs. P14 | | | | | 0.684 | No | 0.311 | No |
| **eAP max rise slope (mV/ms)** | P14 vs. P21 | <0.001 | Yes | <0.001 | Yes | <0.001 | Yes | 0.538 | No |
| | P21 vs. P35 | 0.441 | No | 0.156 | No | 0.507 | No | 0.637 | No |
| | P35 vs. P14 | <0.001 | Yes | <0.001 | Yes | <0.001 | Yes | 0.711 | No |
| **AIS peak amplitude of $d^2V/dt^2$ ($mV/ms^2$)** | P14 vs. P21 | <0.001 | Yes | <0.001 | Yes | <0.001 | Yes | <0.001 | Yes |
| | P21 vs. P35 | 0.519 | No | 0.052 | No | <0.001 | Yes | 0.473 | No |
| | P35 vs. P14 | <0.001 | Yes | <0.001 | Yes | <0.001 | Yes | <0.001 | Yes |
| **Somatic peak amplitude of $d^2V/dt^2$ ($mV/ms^2$)** | P14 vs. P21 | <0.001 | Yes | 0.196 | No | <0.001 | Yes | 0.161 | No |
| | P21 vs. P35 | 0.675 | No | 0.966 | No | 0.047 | Yes | 0.026 | Yes |
| | P35 vs. P14 | <0.001 | Yes | 0.187 | No | 0.007 | Yes | 0.002 | Yes |
| **Delay between the peaks of $d^2V/dt^2$ (ms)** | P14 vs. P21 | <0.001 | Yes | <0.001 | Yes | 0.038 | Yes | <0.001 | Yes |
| | P21 vs. P35 | 0.018 | Yes | 0.043 | Yes | 0.822 | No | <0.001 | Yes |
| | P35 vs. P14 | 0.029 | Yes | <0.001 | Yes | 0.037 | Yes | <0.001 | Yes |
| **AP half-width (ms)** | P14 vs. P21 | <0.001 | Yes | <0.001 | Yes | 0.015 | Yes | 0.005 | Yes |
| | P21 vs. P35 | 0.606 | No | <0.001 | Yes | 0.006 | Yes | 0.023 | Yes |
| | P35 vs. P14 | <0.001 | Yes | <0.001 | Yes | 0.899 | No | 0.426 | No |
| **EPSP half-width (ms)** | P14 vs. P21 | 0.069 | No | <0.001 | Yes | 0.923 | No | 0.122 | No |
| | P21 vs. P35 | 0.892 | No | <0.001 | Yes | 0.069 | No | 0.004 | Yes |
| | P35 vs. P14 | 0.047 | Yes | 0.527 | No | 0.113 | No | 0.292 | No |
| **AP max decay slope (mV/ms)** | P14 vs. P21 | <0.001 | Yes | <0.001 | Yes | <0.001 | Yes | <0.001 | Yes |
| | P21 vs. P35 | 0.006 | Yes | <0.001 | Yes | 0.003 | Yes | 0.433 | No |
| | P35 vs. P14 | <0.001 | Yes | 0.002 | Yes | 0.004 | Yes | 0.005 | Yes |

**Table S1: Additional statistical information regarding Fig. 9.** Two Way ANOVA analysis within genotypes over time.
